# 2,3-bis (phenylamino) quinoxaline - containing compounds display potent activity against Methicillin-resistant *Staphylococcus aureus*, *Enterococcus faecalis* and their biofilms

**DOI:** 10.1101/2024.07.15.603556

**Authors:** Gilda Padalino, Katrina Duggan, Luis A. J. Mur, Jean-Yves Maillard, Andrea Brancale, Karl F. Hoffmann

## Abstract

Antimicrobial resistance remains a global challenge threatening the ability to control diseases caused by bacterial infections. Here, we explored the antimicrobial activity of 2,3-N,N-diphenyl quinoxaline derivatives against representative Gram-positive, Gram-negative and *Mycobacterium* species. Two quinoxaline derivatives (compounds **25** and **31**) demonstrated particularly potent activity against most *Staphylococcus aureus*, *Enterococcus faecium* and *Enterococcus faecalis* strains tested (MIC values between 0.25 to 1 mg/l). These compounds also demonstrated potent antibacterial activity against methicillin-resistant *S. aureus* (MRSA) and vancomycin-resistant *E. faecium/E. faecalis* (VRE) strains. Against an extensive panel of clinically relevant isolates, they further showed comparable or better activity to four currently used antibiotics (vancomycin, teicoplanin, daptomycin and linezolid). Finally, they performed better in preventing *S. aureus* and *E. faecalis* biofilm formation when compared to several other antibiotics. In conclusion, these two quinoxaline derivatives have promising activities that could be further explored as part of efforts to identify urgently needed new antibacterial agents.

---

Antimicrobial resistance (AMR) leads to treatment failure, increased mortality and morbidity as well as spiralling costs for healthcare providers and governments [1]. With an estimated 10 million deaths attributable to AMR by 2050, this public health threat remains a worldwide concern [2]. Recently, an in-depth evaluation from Murray and colleagues showed that 4.95 million deaths were associated with drug-resistant infections globally in 2019 [3]. Among the potential interventions to effectively combat the rise of AMR are those that involve the identification and development of novel antimicrobials [4].

Heterocyclic structures, contained within natural or synthetic products, are increasingly being used as components of new therapeutics. Amongst these, the quinoxaline core represents an important scaffold associated with many biologically- and pharmacologically-active properties useful for treating both non-communicable diseases and infectious agents [5, 6]. For example, quinoxaline derivatives possess potent anti-parasitic activities against *Leishmania* [7], *Trypanosoma* [8], *Plasmodium* [9] and *Schistosoma* [10] species.

During our search for broadly-active anthelmintics to control the neglected tropical disease Schistosomiasis, we recently designed, synthesised and evaluated a small library of quinoxaline analogues against *Schistosoma mansoni*, *Schistosoma japonicum* and *Schistosoma haematobium* [11]. While these compounds demonstrated anti-schistosomal potencies at nanomolar concentrations, they also displayed structural similarities to previously-described, antibacterial, quinoxaline-containing compounds [12].

With the increasing concerns around the emergence of resistance [13, 14] and tolerance [15, 16] to currently used antibiotics, we decided to further investigate the broader antibacterial potential of this family of quinoxaline-containing compounds. Here, we first tested a small number of 2,3-*N*,*N*-diphenyl quinoxaline derivatives against a wide panel of bacterial strains (Gram-positive bacteria, Gram-negative bacteria and *Mycobacterium smegmatis*) to gather preliminary information about structure–activity relationship (SAR). We subsequently progressed more detailed antibacterial screens with selected compounds against defined strains of clinical relevance, particularly against antibiotic-resistant isolates.

We used both a standard microdilution broth assay to determine minimal inhibitory concentrations (MIC) and a biofilm test to measure minimal biofilm eradication concentrations (MBEC). Indeed bacterial biofilms are less susceptible to antibiotics [17] and are clinically more relevant [17, 18].

By doing so, our results demonstrate the relevance of two of our synthesised quinoxaline derivatives (compound **25** and compound **31**) against a range of bacteria, warranting further investigations.

## Results

### Determination of *in vitro* antibacterial activity against ATCC/NTCT isolates (Phase 1)

We recently reported the identification of quinoxaline derivatives as part of a high-throughput *ex vivo* screening campaign to identify potent anti-schistosomal compounds [11]. Medicinal chemistry optimisation resulted in the generation of 5 *N*-aryl analogues (compounds **25**, **30-32** and **35**) and two *N*-phenyl-alkyl analogues (compounds **36-37**) created via a one-step reaction using 2,3-Dichloro-6-nitroquinoxaline as the starting material (referred here as 2Cl-Q - **Scheme 1**). The 6-acyl derivatives were obtained from a classical catalytic hydrogenation of the 6-nitro-substituted quinoxaline **22** into the amino derivative, before a final acylation to assemble the analogues **22c**-**22g** (**Scheme 1**). Due to solubility restrictions, only 13 compounds (amongst the originally 21 synthesised in [11]) were selected for antibacterial screening (compounds **25, 30-32**, **35-37, 22b-22g** as well as the 2,3-Dichloro-6-nitroquinoxaline (**2Cl-Q**) and its 6-nitroquinoxaline-2,3-diol derivative starting materials, **Scheme 1** - adapted from [11]).

**Scheme 1.**
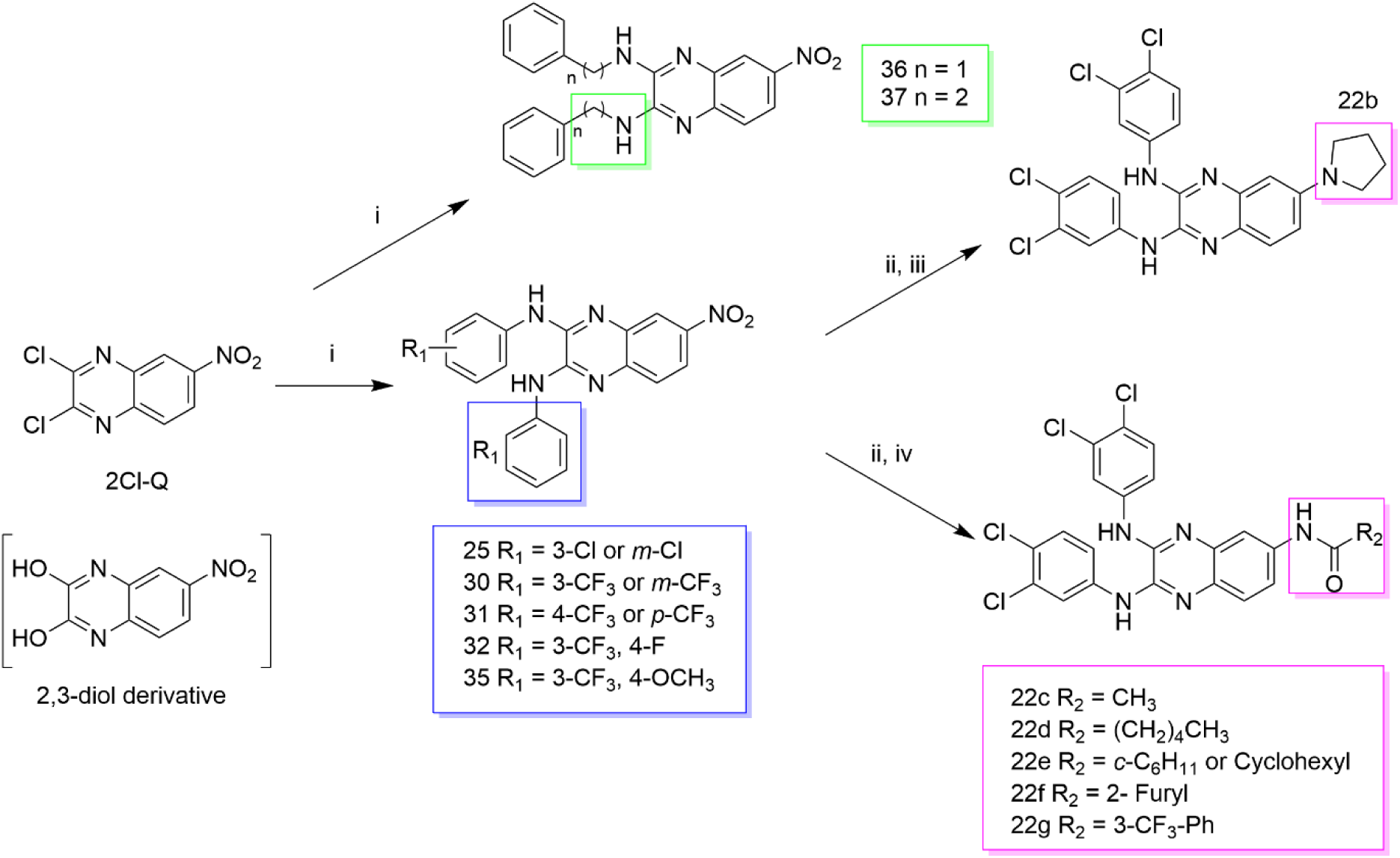
Synthesis of the 2,3-bis(phenylamino)-quinoxaline series. The synthesised derivatives are grouped by structural similarity: the *N*-aryl analogues are highlighted with a blue box, the *N*-phenyl-alkyl analogues are contained in the green box and the 6-acyl derivatives are included in a magenta box. Reagents and conditions: (i) different substituted anilines (compounds **22**–**35**) or phenyl-alkyl amines (**36** and **37**), anhydrous DMSO, 130 °C, 30 min; (ii) H_2_, cat. Pd/C, AcOEt, rt, 2h; (iii) Br(CH_2_)_2_Br, K_2_CO_3_, CH_3_CN, MWI (300 W), 150 °C, 15 min; (iv) R_2_COCl, anhydrous Pyr, anhydrous DCM, 0 °C → rt, 1 h. This figure is adapted from [11] with particular focus only on the quinoxaline derivatives (**Table S1A**) and bacterial species (**Table S1B**) investigated in this study.

The minimum inhibitory concentration (MIC) of this library was initially assessed by the broth microdilution method [19]. Representative species derived from Gram-positive and Gram-negative strains as well as *M. smegmatis* were first screened with all compounds at two concentrations (125.00 and 62.50 mg/l). Compounds showing inhibition of bacterial growth at 62.50 mg/l were further tested in a secondary dose response assay. From these screens, the minimal concentrations of 15 quinoxaline analogues that reduced visible growth were calculated for each bacteria strain (**Table 1**).

**Table 1.** Minimum Inhibitory Concentrations (MICs, mg/l) of 15 quinoxaline derivatives.

|  | MIC (mg/l) |  |  |  |  |  |
| --- | --- | --- | --- | --- | --- | --- |
|  | Gram-positive |  |  | Gram-negative |  | Mycobacteria* |
| Entry | *S. aureus* | *S. epidermidis* | *B. cereus* | *E. coli* | *P. aeruginosa* | *M. smegmatis* mc<sup>2</sup>15 |
| 2,3-dichloro-6-nitroquinoxaline/<br>2Cl-Q | <3.91 | <62.50 | <62.50 | 62.50 | 125 | 1.95 |
| 6-nitroquinoxaline-2,3-diol | 125 | - | - | >125 | - | >125 |
| 25 | <3.91 | <62.50 | 125 | >125 | >125 | >125 |
| 30 | 7.81 | >125 | >125 | >125 | >125 | >125 |
| 31 | 0.98 | <62.50 | <62.50 | >125 | >125 | 31.25 |
| 32 | 1.95 | <62.50 | 125 | 62.5 | >125 | 62.50 |
| 35 | 62.50 | <62.50 | <62.50 | >125 | >125 | >125 |
| 36 | >125 | - | - | >125 | - | >125 |
| 37 | >125 | - | - | >125 | - | 31.25 |
| 22b | 15.63 | >125 | >125 | >125 | >125 | >125 |
| 22c | 62.50 | <62.50 | <62.50 | >125 | >125 | >125 |
| 22d | 62.50 | >125 | 125 | >125 | >125 | >125 |
| 22e | >125 | - | - | >125 | - | 125 |
| 22f | 31.25 | 125 | <62.50 | >125 | >125 | >125 |
| 22g | >125 | - | - | >125 | - | 125 |

\* 48 hours incubation with drug. Other bacteria strains were incubated for 24 hours.
Strains used: *S. aureus* - ATCC 29213; *Staphylococcus epidermidis* - NTCT11077; *Bacillus cereus* - ATCC 14579; *Escherichia coli* - ATCC 25922; *Pseudomonas aeruginosa* ATCC 27853, *M. smegmatis* mc<sup>2</sup>15 ATCC 700084. Concentration tested: 125, 62.50, 31.25, 15.63, 7.81, 3.91, 1.95, 0.98 µg/ml; Green cells indicate MIC values of 31.25 mg/l or lower.
Cells containing the value '>125' indicate that bacterial growth was still observed at the highest concentration tested (125 mg/l). Cells containing the value '<62.5' or '< 3.91' indicate that there was no visible bacterial growth at that concentration (62.5 and 3.91 mg/l, respectively). A further titration (below 62.50 or 3.91 mg/l,
respectively) would be needed to define the exact value of MIC. Cells containing ‘-’ indicated that the MIC of that compound was not determined.

Of the six bacteria species tested, *Escherichia coli*, *Pseudomonas aeruginosa, Bacillus cereus* and *Staphylococcus epidermidis* were minimally affected (MIC > 62.5 - 125 mg/l) by the compounds. Two compounds (**31** and **37**) demonstrated some antimycobacterial activity, with the initial building block (2,3-Dichloro-6-nitroquinoxaline) being the most active (MIC 1.95 mg/l). In contrast, *Staphylococcus aureus* showed the highest susceptibility to these compounds with **31** and **32** producing MICs of 0.98 and 1.95 mg/l, respectively. Only 5 of the 15 compounds tested (6-nitroquinoxaline-2,3-diol, **36**, **37**, **22e** and **22g**) were not effective against this bacterium (MIC ≥ 125 mg/l). Compound **25** and 2,3-dichloro-6-nitroquinoxaline showed some activity with a MIC below 3.91 mg/l.

The antibacterial properties of 7 of the most active compounds against *S. aureus* prompted further studies on three methicillin-resistant *S. aureus* (MRSA) strains (**Table 2**).

**Table 2.** Minimum Inhibitory Concentrations (MICs, mg/l) against three selected MRSA strains compared to a reference *S. aureus* strain.

| Entry | MIC (mg/l) |  |  |  |
| --- | --- | --- | --- | --- |
|  | Reference *S. aureus* * | MRSA |  |  |
|  | ATCC 29213 | USA 300 | ATCC 33591 | EM RSA |
| 2,3-dichloro-6-nitroquinoxaline/<br>2Cl-Q | <3.91 | 15.63 | 15.63 | 15.63 |
| 6-nitroquinoxaline-2,3-diol | 125 | - | - | - |
| 25 | <3.91 | 3.91 | 3.91 | 3.91 |
| 30 | 7.81 | >125 | >125 | >125 |
| 31 | 0.98 | <3.91 | <3.91 | 3.91 |
| 32 | 1.95 | <3.91 | <3.91 | 7.81 |
| 35 | 62.50 | 15.63 | 31.25 | 31.25 |
| 36 | >125 | - | - | - |
| 37 | >125 | - | - | - |
| 22b | 15.63 | 62.50 | 31.25 | 62.50 |
| 22c | 62.50 | - | - | - |
| 22d | 62.50 | - | - | - |
| 22e | >125 | - | - | - |
| 22f | 31.25 | 62.50 | 15.63 | 62.50 |
| 22g | >125 | - | - | - |

\*MICs values against *S. aureus* reported here as reference (from **Table 1**). Green cells indicate MIC values of 31.25 mg/l or lower. Those compounds were selected for further screens against MRSA strains. Strains used: standard *S. aureus* - ATCC 29213; Methicillin-resistant *S. aureus* (MRSA) isolates - USA300, ATCC 33591, EM RSA.

Here, some compounds (e.g., 2,3-dichloro-6-nitroquinoxaline and compounds **30, 22b** and **22g**) lost activity against some or all of the MRSA strains tested. Compounds **25**, **31** and **32** retained some activity on MRSA strains, but they were not as potent when compared to the *S. aureus* reference strain. Compound **35** was the only compound that showed an increased potency against the MRSA strains (15.53-31.25 mg/l compared to a MIC of 62.50 mg/l for the standard *S. aureus* strain).

### Determination of *in vitro* antibacterial activity against clinically relevant bacteria strains (Phases 2 and 3)

Based on compound **25**, **31**, **32**, **22f**, **35** and 2,3-dichloro-6-nitroquinoxaline/**2Cl-Q**’s activities against MRSA strains, a wider antibacterial screen (**Figure 1**) was conducted on 19 clinically relevant bacterial species/strains (**Table S2A**).

**Figure 1.**
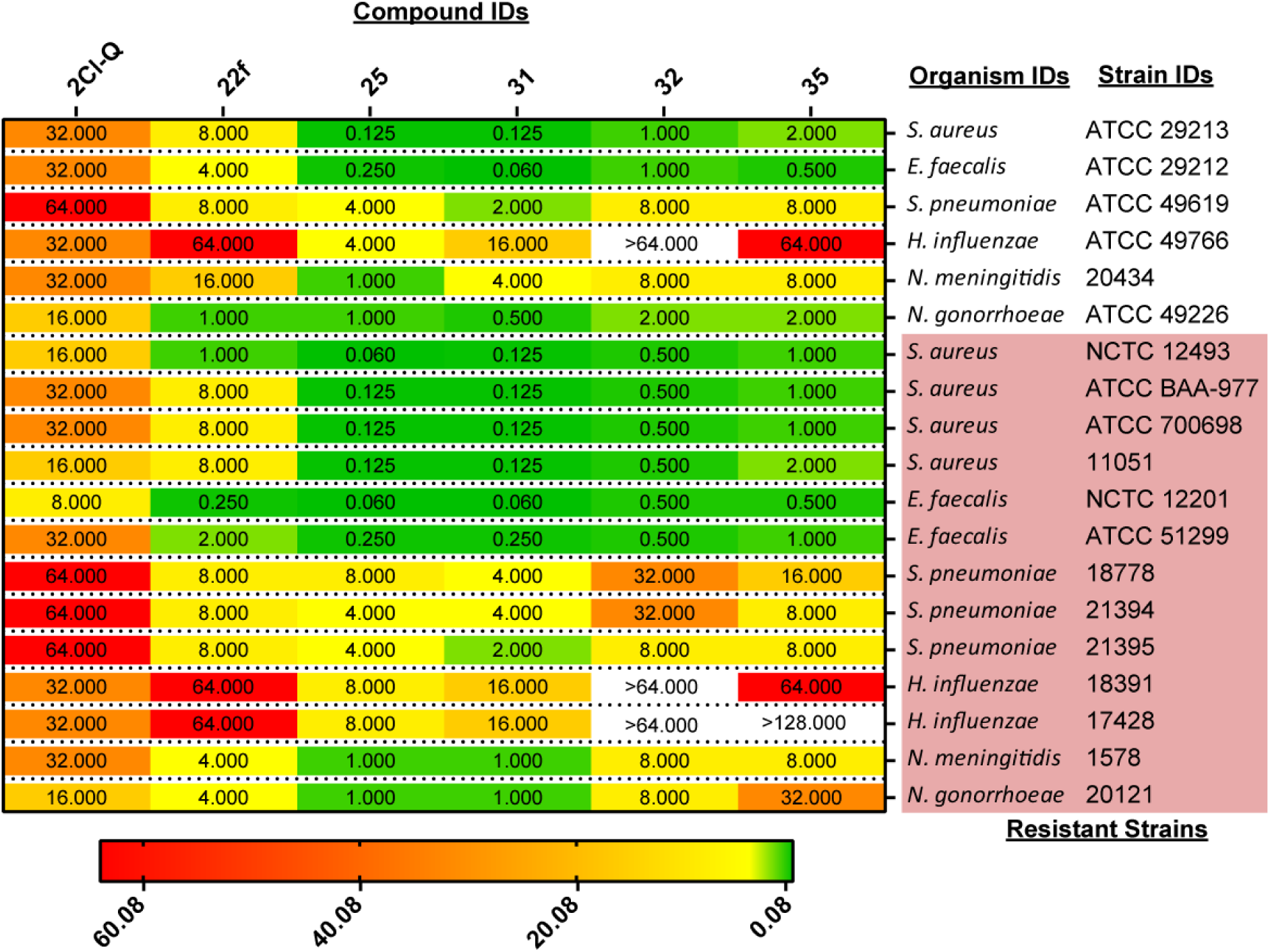
Minimum Inhibitory Concentrations (MICs, mg/l) against 19 clinically relevant strains. More information about the strains can be found in **Table S2A.** MICs (expressed in mg/l) are indicated. A colour code was used based on the highest (64 mg/l – in red) to the lowest value (0.06 mg/l – in green) contained within the dataset (coloured bar, in mg/l, is at the bottom of the picture). Relevant strains with resistance mechanisms are highlighted in pink – more details on the specific resistance and mechanism can be found in **Table S2A.** The complete antibacterial dataset is available in **Table S2B.** 2Cl-Q = 2,3-dichloro-6-nitroquinoxaline.

Amongst the compounds tested, **2Cl-Q** showed minimal activities (MICs ≥ 8 mg/l) against the selected strains. Compound **22f** displayed activities against *Neisseria gonorrhoeae* (ATCC 49226; MIC of 1 mg/l), one flucloxacillin-resistant *S. aureus* strain (*S. aureus* NCTC 12493; MIC = 1 mg/l) and two vancomycin-resistant *Enterococcus faecalis* (VRE) strains (NCTC 12201 - MIC = 0.250 mg/l and ATCC 51299 - MIC = 2 mg/l). Compounds **25**, **31**, **32** and **35** were active against *S. aureus* and *E. faecalis* wild type strains. However, these compounds were even more active against the drug-resistant *S. aureus* and *E. faecalis* strains with compound **25** and **31** generally being more active than the other two (**Figure 1**).

Exploring the wider panel of Gram-positive bacteria strains, compound **31** was the only compound with activity (MIC = 2 mg/l) against *Streptococcus pneumoniae* (ATCC 49619), the most common cause of community-acquired pneumonia and one of nine bacteria of international concern (World Health Organization; WHO [29]). This compound also retained its activity against the MLS-resistant *S. pneumoniae* strain (21395, MIC = 2 mg/l), but was less active against 18778 and 21394 (MIC = 4 mg/l).

Focusing on the Gram-negative species, compounds **25** and **31** were also the only two compounds active against *Neisseria* species and *Haemophilus influenzae*. In terms of *H. influenzae*, compound **25** demonstrated greater activity (compared to **31**) against both standard-(ATCC 49766, MIC = 4 and 16 mg/l for compounds **25** and **31**, respectively) and resistant-strains (18391, 17428 with MIC = 8 and 16 mg/l for compounds **25** and **31**, respectively).

Compounds **31** and **25** showed the broadest and most potent antibacterial activities against all species/strains investigated, including the thirteen drug resistant strains (**Figure 1**). Therefore, these two quinoxalines were selected for further investigations against additional drug-resistant strains of both *S. aureus* and *E. faecalis* (**Table 3**).

**Table 3.** Minimum Inhibitory Concentrations (MICs, mg/l) against drug-resistant *S. aureus* and *E. faecalis* strains.

| Organism ID | Reference strain | Resistance | Mechanism | Compound IDs |  |
| --- | --- | --- | --- | --- | --- |
|  |  |  |  | 25<br>(MICs, mg/l) | 31<br>(MICs, mg/l) |
| *E. faecalis* | ATCC 29212 | - | - | 0.250 | 0.060 |
| *E. faecalis* | NCTC 12201 | Vancomycin | *vanA* | 0.060 | 0.060 |
| *E. faecalis* | ATCC 51299 | Vancomycin | *vanB* | 0.250 | 0.250 |
| *S. aureus* | ATCC 29213 | - | - | 0.125 | 0.125 |
| *S. aureus* | NCTC 12493 | Flucloxacillin | *mecA* | 0.060 | 0.125 |
| *S. aureus* | ATCC BAA-977 | ERY/CLIND | MLSB | 0.125 | 0.125 |
| *S. aureus* | ATCC 700698 | Vancomycin | hVISA | 0.125 | 0.125 |
| *S. aureus* | 11051 | Tetracycline | - | 0.125 | 0.125 |

Both compounds were particularly active against vancomycin-resistant strains of *E. faecalis*, especially against the NCTC 12201 strain (MIC = 0.60 mg/l). They additionally demonstrated equivalent potencies against the flucloxacillin-, vancomycin-, tetracycline- and erythromycin/clindamycin-resistant *S. aureus* strains (**Table 3**).

### Activity of compounds 25 and 31 on additional clinically relevant bacterial strains (Phase 3)

Based on the promising antibacterial effects of compounds **25** and **31** on representative Gram-positive bacteria of clinical importance, a more expansive panel of Gram-positive bacterial strains was next subjected to MIC investigations (**Figure 2** and **Table S3**). In addition to *S. aureus* (**Figure 2A**) and *E. faecalis* (**Figure 2B**), *Enterococcus faecium* (**Figure 2C**) was also included in these assays due to broader resistance and higher virulence than *E. faecalis* [30].

**Figure 2.**
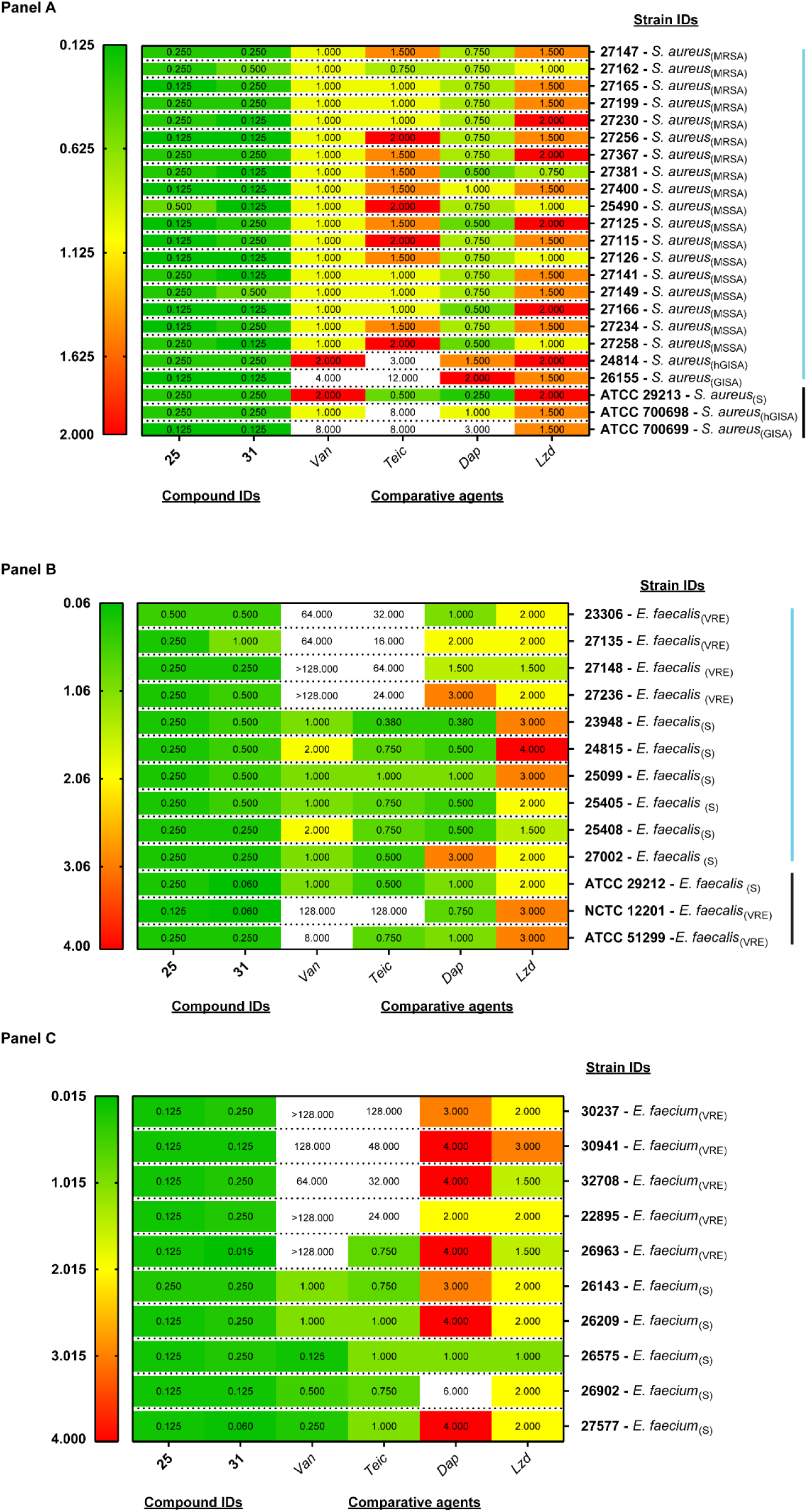
Minimum Inhibitory Concentrations (MICs, mg/l) of two selected compounds and four comparative agents against a panel of *S. aureus* (Panel A), *E. faecalis* (Panel B) and *E. faecium* (Panel C). **Panel A** - All clinical strains of *S. aureus* (blue line), except for the final three standard *S. aureus* strains (ATCC 29213, ATCC 700698 and ATCC 700699, black line). MSSA = methicillin-sensitive *Staphylococcus aureus*; S = sensitive; GISA = glycopeptide-intermediate *S. aureus*; hGISA = hetero-GISA; **Panel B** - All clinical strains of *E. faecalis* (blue line), except for the final three ATCC/NTC strains (black line). VRE = vancomycin-resistant enterococci; S = sensitive short for MSSA - methicillin-sensitive *S. aureus*; **Panel C** - VRE = vancomycin-resistant enterococci; S = sensitive. Comparative agents: *Van* – Vancomycin; *Lzd* – Linezolid; *Teic* - Teicoplanin; *Dap* – Daptomycin. A colour code was used based on the highest (in red) to the lowest MIC value (in green) of the dataset (coloured bar, mg/l, is included at the left-hand side of the picture). The highest range of the colour mapping was set to 2.00 (**Panel A**) or 4.00 (**Panels B** and **C**) for visualisation purposes.

Vancomycin, teicoplanin, linezolid and daptomycin MICs were first compared to the two quinoxaline analogues (**Table S3A** and **Table S3B**). Regarding *S. aureus*, three control strains (methicillin-sensitive *Staphylococcus aureus* strain ATCC 29213 - MSSA - and two MRSA strains ATCC 700698 and 700699) were evaluated along with twenty clinical isolates. Compounds **25** and **31** showed a good antibacterial profile across all *S. aureus* isolates under analysis with MICs ranging from 0.125 to 0.500 mg/l (**Figure 2A**).

A heterogeneous MIC spectrum for the four clinical antibiotics was identified for the *S. aureus* standard stains and clinical isolates with linezolid being the least potent. Both quinoxaline analogues demonstrated greater potencies than vancomycin, teicoplanin and linezolid with comparable activity to daptomycin in some cases. When looking at the GISA strains, the two test compounds outperformed the first line treatment (vancomycin and teicoplanin - **Figure 2A**).

Compounds **25** and **31** were next tested against a panel of 13 *E. faecalis* strains (**Figure 2B**); these included three standard strains (sensitive strain ATCC 29212 and two vancomycin-resistant enterococci (VRE) strains (NTCC 12201 and ATCC 51299)) and ten clinically relevant strains (**Table S3A**). Both compounds **25** and **31** showed antibacterial activity against all *Enterococci* tested. While compound **25** appeared equally active across the strains under analysis, compound **31** had a much wider range of MICs from 0.060 mg/l against ATCC 29212 and NCTC 12201 to 1 mg/l against clinical isolate 27135.

Lastly, the antibacterial activities of compounds **25** and **31** were investigated against a selection of clinically relevant *E. faecium* strains (vancomycin-sensitive or resistant exemplars; **Figure 2C**). Both compounds had MIC values lower than 0.250 mg/ml against all strains examined (both sensitive and resistant strains of standard and clinical isolates of *E. faecium*). This represents a better antimicrobial profile over the second line agents (daptomycin and linezolid) and, in some cases, an equal (or even better activity) when compared to first line agents (vancomycin and teicoplanin).

### Determination of MIC following pre-exposure to quinoxaline analogues 25 and 31 (Phase 4)

To determine if pre-exposure to compounds **25** and **31** altered antimicrobial susceptibility, *S. aureus* and *E. faecalis* reference strains were exposed to sub-MIC levels (50% of the MIC for 16-24 hrs) of these quinoxaline analogues. MICs of pre-exposed (Spe) strains to controls without pre-exposure (Sc) were subsequently compared (**Table 4**). Following pre-exposure of bacteria to sub-MIC levels of compound **25**, MICs increased between 2 and 8-fold. Pre-exposure of bacteria to sub-MIC levels of compound **31** increased MICs to between 64- and 256-fold.

**Table 4.**
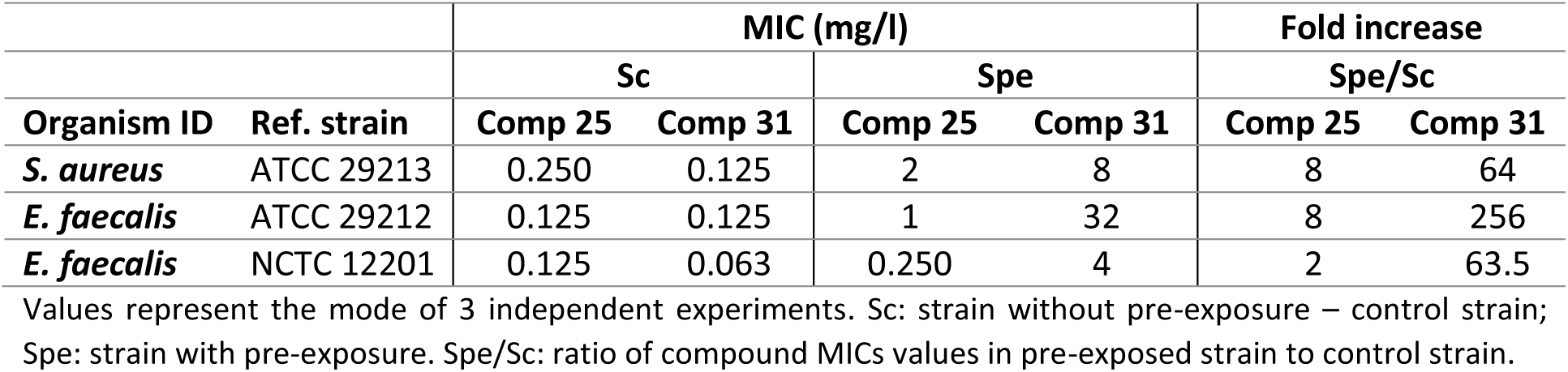
Pre-exposure of *S. aureus* and *E. faecalis* to quinoxaline analogues decreases their sensitivities.

Since pre-exposure of bacteria to the two quinoxaline analogues decreased their susceptibility, MICs of commonly used antibiotics (ampicillin, imipenem, vancomycin, levofloxacin, ciprofloxacin, trimethoprim/sulfamethoxazole, cefotixin, gentamicin, erythromycin, tetracycline, rifampicin and benzylpenicillin) were subsequently determined for *S. aureus* and *E. faecalis* with and without pre-exposure to compounds **25** and **31** (**Table 5**).

**Table 5.** Pre-exposure of *S. aureus* and *E. faecalis* to compounds 25 and 31 and impact on cross resistance to clinically available antibiotics.

|  | *S. aureus* ATTC 29213 |  |  | *E. faecalis* NCTC 12201 |  |  | *E. faecalis* ATCC 29212 |  |  |
| --- | --- | --- | --- | --- | --- | --- | --- | --- | --- |
|  | Sc | Spe<sub>25</sub> | Spe<sub>31</sub> | Sc | Spe<sub>25</sub> | Spe<sub>31</sub> | Sc | Spe<sub>25</sub> | Spe<sub>31</sub> |
| **Ampicillin** | ND | ND | ND | 19.8 | 19.0 | 20.0 | 17.2 | 17.8 | 17.9 |
| **Imipenem** | ND | ND | ND | 26.4 | 26.5 | 26.5 | 27.1 | 27.8 | 27.8 |
| **Vancomycin** | ND | ND | ND | 10.7 | 10.9 | 15.3 | 15.9 | 16.6 | 15.6 |
| **Levofloxacin** | ND | ND | ND | 20.4 | 21.2 | 21.5 | 22.6 | 22.0 | 23.0 |
| **Ciprofloxacin** | 22.0 | 21.9 | 22.0 | 21.6 | 21.9 | 22.3 | 24.0 | 24.0 | 24. |
| **Tri/Sulf** | 29.9 | 31.9 | 30.0 | 28.6 | 28.7 | 0 | 32.0 | 31.5 | 32.8 |
| **Cefotixin** | 28 | 28.2 | 28.3 | ND | ND | ND | ND | ND | ND |
| **Gentamicin** | 21.5 | 22.4 | 22.2 | ND | ND | ND | ND | ND | ND |
| **Erythromycin** | 25.7 | 27.2 | 26.6 | ND | ND | ND | ND | ND | ND |
| **Tetracycline** | 24.8 | 24.8 | 25.8 | ND | ND | ND | ND | ND | ND |
| **Rifampicin** | 31.6 | 32.2 | 33.0 | ND | ND | ND | ND | ND | ND |
| **Benzylpenicillin** | 14.8 | 14.2 | 13.0 | ND | ND | ND | ND | ND | ND |

Zones of inhibition (mm) produced by antibiotic loaded disks against *S. aureus* and *E. faecalis* species. Sc: strain without exposure; Spe<sub>25</sub>: strain with exposure to compound **25**; Spe<sub>31</sub>: strain with exposure to compound **31**; Tri/Sulf: Trimethoprim/sulfamethoxazole; ND: Not Determined; Bacteria were deemed susceptible (in green) or resistant (in red) according to EUCAST clinical breakpoints.

Pre-exposure of *S. aureus* ATTC 29213 or *E. faecalis* ATCC 29212 to sub-inhibitory concentrations of either compound did not alter these strains’ susceptibility to any of the antibiotics tested. In contrast, pre-exposure of *E. faecalis* NCTC 12201 to compound **31** (but not compound **25**) led to vancomycin susceptibility with a zone of inhibition greater than the clinical breakpoint. Additionally, *E. faecalis* NCTC 12201 pre-exposed to a sub-inhibitory concentration of compound **31** (but not compound **25**) altered trimethoprim-susceptibility to a trimethoprim-resistant phenotype (**Table 5**).

### Minimum biocidal concentration (MBC) determination and scanning electron microscopy (SEM) analyses

To complement the extensive MIC testing of compounds **25** and **31**, minimum biocidal concentrations of these quinoxaline analogues were also determined for both *S. aureus* and *E. faecalis* (**Table 6**). Compound **25** demonstrated greater bactericidal activity against both *S. aureus* and *E. faecalis* when compared to compound **31** (**Table 6**).

**Table 6.** MBC values of compounds 25 and 31 against *S. aureus* and *E. faecalis*.

| Organism ID | Reference strain | MBC (mg/l) |  |
| --- | --- | --- | --- |
|  |  | Comp 25 | Comp 31 |
| *S. aureus* | ATCC 29213 | 1 | 8 |
| *E. faecalis* | ATCC 29212 | 4 | 8 |
| *E. faecalis* | NCTC 12201 | 4 | 8 |

Values represent the mode of 3 independent experiments.

SEM images of both *S. aureus* and *E. faecalis* exposed to double the MBC of compounds **25** and **31** revealed the gross structural damage in comparison to untreated cells (**Figure 3**, Panels **A**, **B**, **G**, **H**). The morphology of *S. aureus* exposed to compound **25** changed from round, well defined cells in the untreated cultures (**Panels A** and **B**) to dimpled and irregular cells (**Panels C** and **D**). The damage to *S. aureus* cells treated with compound **31** appeared even more pronounced, with greater levels of structural damage and possible loss of intracellular contents (**Panels E** and **F**). Similarly, *E. faecalis* exposed to compound **25** (**Panels I** and **J**) and **31** (**Panels K** and **L**) showed severe structural damage, including wrinkled cell surfaces and cells that appear to have collapsed.

**Figure 3.**
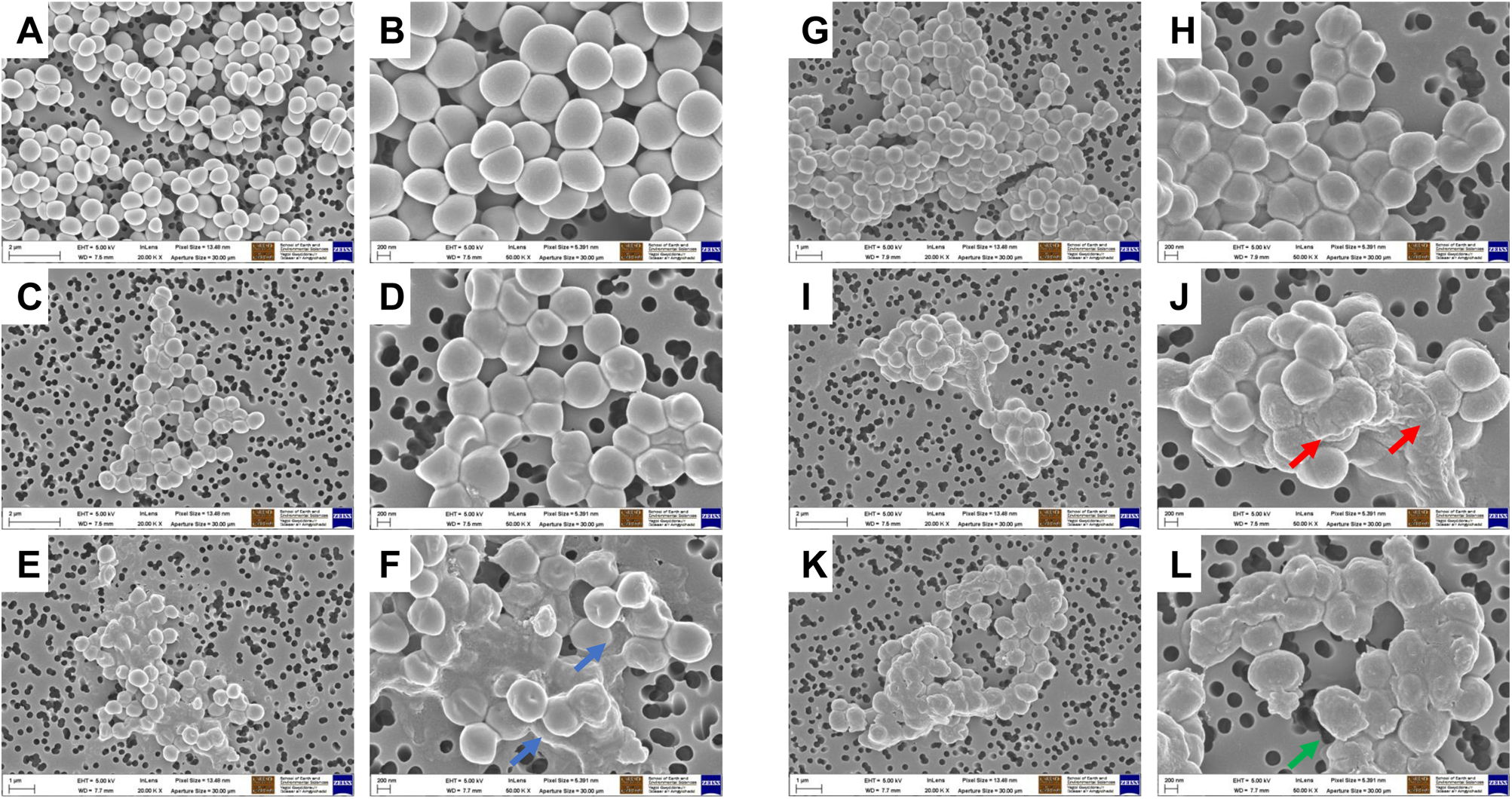
Quinoxaline analogues induce phenotypic alterations to the surface of both *S. aureus* and *E. faecalis*. Scanning electron micrographs of *S. aureus* ATCC 29213 (**A** – **F**) and *E. faecalis* NCTC 12201 (**G** – **L**). **Panels A** and **G** and **Panels B** and **H** represent untreated cells captured at 20,000- and 50,000-times magnification, respectively. Bacteria treated with 2x MBC concentration of compound **25** are depicted in **Panels C** and **I** as well as **Panels D** and **J** at 20,000- and 50,000-times magnification, respectively. Bacteria treated with 2 x MBC concentration of compound **31** are shown in **Panels E** and **K** as well as **Panels F** and **L** at 20,000- and 50,000-times magnification, respectively. Blue arrows: loss of cellular content; red arrow: wrinkled cell surfaces; green arrow: cells that appear to have collapsed

### Estimation of Minimum Biofilm Eradication Concentration (MBEC)

The efficacy of compounds **25** and **31** to eradicate pre-formed biofilms of *S. aureus* and *E. faecalis* was next determined and compared to a group of known antibiotics (vancomycin, rifampicin, linezolid, teicoplanin and sparfloxacin) (**Table 7**). A MBEC of 256 mg/l was found for both quinoxaline analogues against *S. aureus* ATCC 29213 and *E. faecalis* ATCC 29212 strains. While compound **25** also yielded a MBEC of 256 mg/L against *E. faecalis* ATCC 12201, compound **31** was less effective in eradicating biofilms caused by this bacterium. Except for rifampicin against *S. aureus* (MBEC < 128 mg/l), the other four antibiotics (vancomycin, linezolid, teicoplanin and sparfloxacin) all had higher MBECs (>512 or > 1024 mg/L) than compounds **25** and **31** (**Table 7**).

**Table 7.** Quinoxaline analogues showed better anti-biofilm activities (MBEC) compared to five selected comparative antibiotics.

|  | MBEC (mg/l) |  |  |
| --- | --- | --- | --- |
|  | *S. aureus*<br>ATCC 29213 | *E. faecalis*<br>ATCC 29212 | *E. faecalis*<br>NCTC 12201 |
| **Compound 25** | 256 | 256 | 256 |
| **Compound 31** | 256 | 256 | >256 |
| **Vancomycin** | >512 | >512 | >512 |
| **Rifampicin** | <128 | >512 | >512 |
| **Linezolid** | > 1024 | > 1024 | > 1024 |
| **Teicoplanin** | > 1024 | > 1024 | > 1024 |
| **Sparfloxacin** | > 1024 | > 1024 | > 1024 |

Values represent the mode of 3 independent experiments.

## Discussion

### Structure-activity relationship (SAR) studies based on Phase 1 results

The antibacterial data collected for a small number of quinoxaline analogues in Phase 1 of this study highlighted that the central quinoxaline scaffold was responsible for some activity, particularly against *S. aureus* (2,3-dichloro-6-nitroquinoxaline, **Table 1**). The hydroxy derivative (6-nitroquinoxaline-2,3-diol), in contrast, had poor activity against *S. aureus* and was totally ineffective against Gram-negative bacteria and mycobacteria.

Although limited number of compounds were screened, preliminary structural activity relationship (SAR) analyses highlighted the importance of aromatic ring substitutions and the functionalisation of position 6 of the central scaffold (highlighted in blue and magenta, respectively in **Scheme 1**) for the antimicrobial activity. The *N*-aromatic derivatives of the central quinoxaline scaffold showed antibacterial activity primarily against Gram-positive bacteria with compounds **25**, **31** and **32** being particularly potent (**Table 8**). Regarding the effect of *para* or *meta* substitution, the para-trifluoromethyl (compound **31**) resulted in an increased activity against both *S. aureus* and *M. smegmatis* when compared to the meta isomer (compound **30**). The introduction of a fluorine (compound **32**) led to increased antibacterial activity when compared to the parent compound (the meta-trifluoromethyl derivative **30**) as previously observed with other antibacterial small molecules [31]. However, the combination of a methoxy substituent with a trifluoromethyl group (compound **35**) led to a decrease in activity (against *S. aureus*) when compared to compound **30**.

**Table 8.**
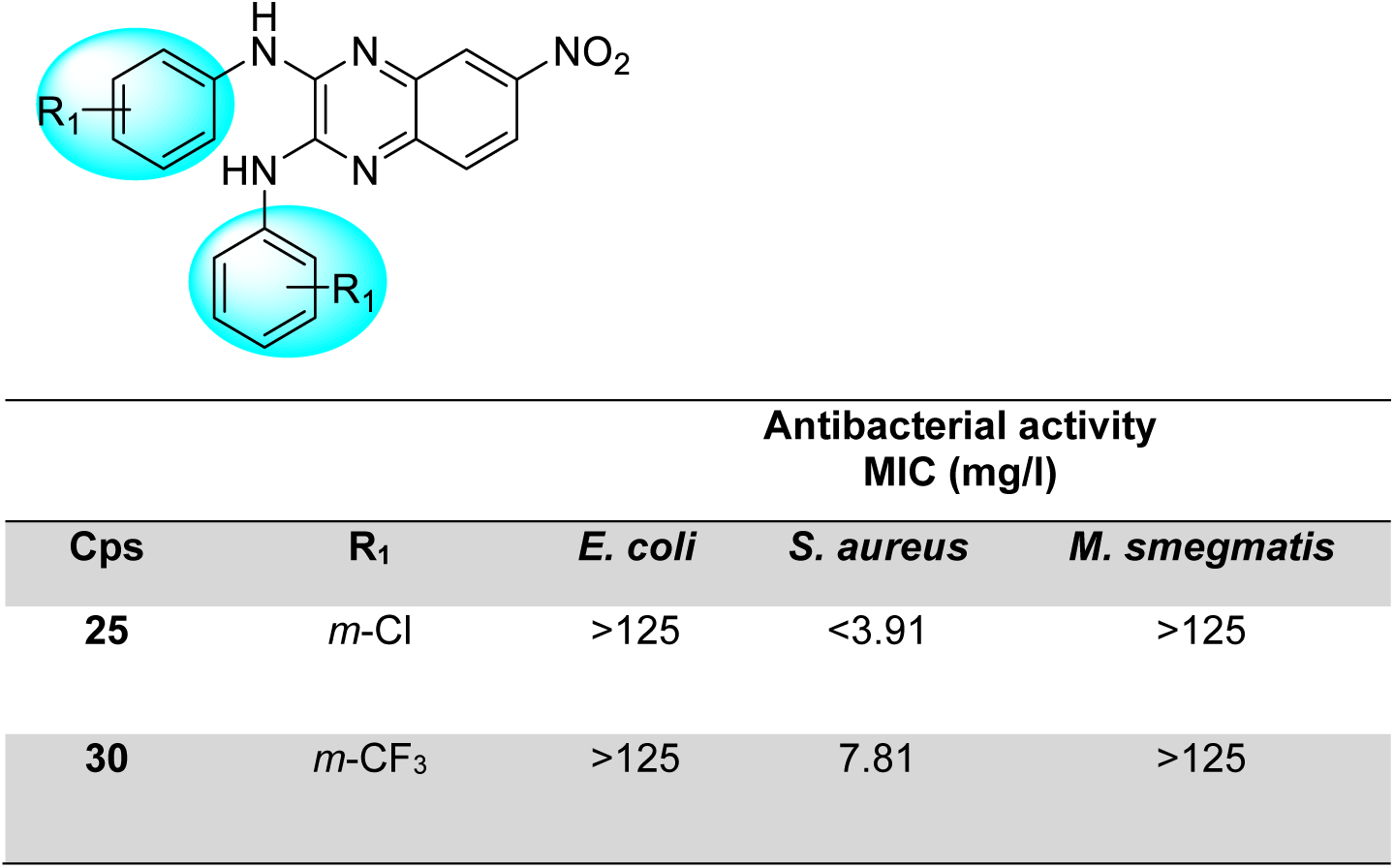
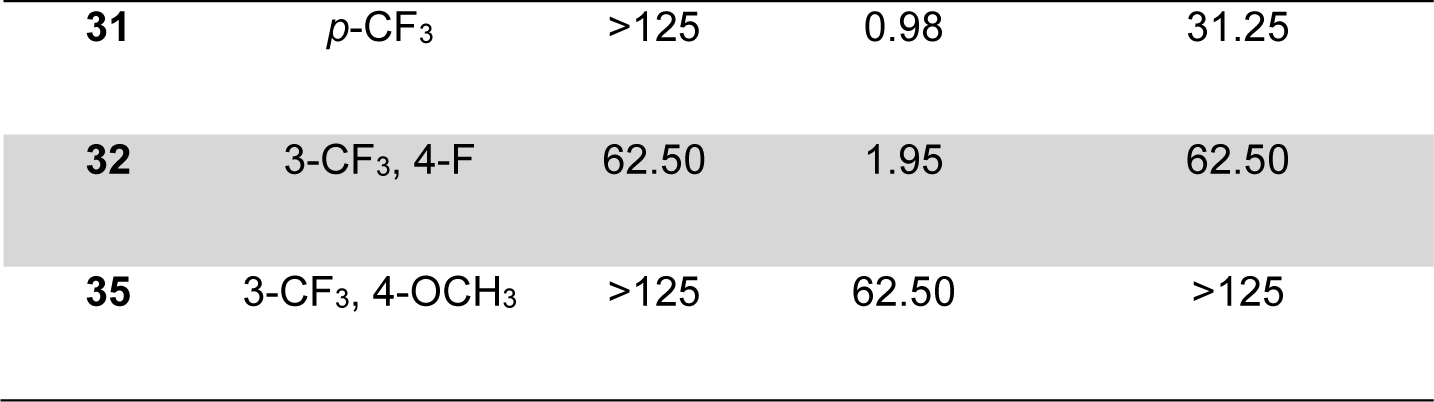
Antibacterial activity of the *N*-aromatic quinoxaline analogues. (R_1_: residue of aromatic ring)

| Antibacterial activity<br>MIC (mg/l) |  |  |  |  |
| --- | --- | --- | --- | --- |
| Cps | R<sub>1</sub> | *E. coli* | *S. aureus* | *M. smegmatis* |
| 25 | *m*-Cl | >125 | <3.91 | >125 |
| 30 | *m*-CF<sub>3</sub> | >125 | 7.81 | >125 |
| **31** | *p*-CF<sub>3</sub> | >125 | 0.98 | 31.25 |
| **32** | 3-CF<sub>3</sub>, 4-F | 62.50 | 1.95 | 62.50 |
| **35** | 3-CF<sub>3</sub>, 4-OCH<sub>3</sub> | >125 | 62.50 | >125 |

One of the original medicinal chemistry optimisations performed on these quinoxaline derivatives involved modifying the C-6 nitro group in order to mitigate cytotoxicity [11]. Therefore, the antibacterial activity of these C-6 derivatives was next explored in order to gain preliminary information about their SARs (**Table 9**). Overall, these analogues showed low activity on the selected bacterial strains suggesting that the nitro group on the C-6 position might be essential for their antibacterial effects, a finding similar to what has previously been reported [32, 33]. The preliminary investigation of these quinoxaline derivatives showed low or no activity at all against *E. coli* and mycobacteria supporting the idea that a lipid rich barrier (outer lipopolysaccharide membrane in Gram-negative or free lipid and mycolate layer in mycobacteria) is an impediment to the uptake of these compounds by these bacteria [34, 35]. While *S. aureus* was somewhat affected by the C-6 nitro-containing compound **22b** (MIC = 15.63 mg/l), the other three compounds lacking a C-6 nitro group (**22c**, **22d** and **22f**) were much less active. These findings broadly suggest that the nitro group offers less versatility for the antibacterial activity compared to the *N*-aromatic rings with further SAR conclusions related to the antibacterial activity of quinoxaline-containing analogues of this study summarised in **Figure 4**.

**Table 9.**
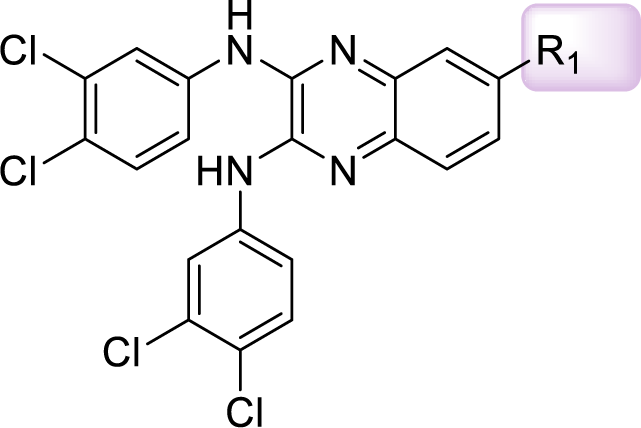
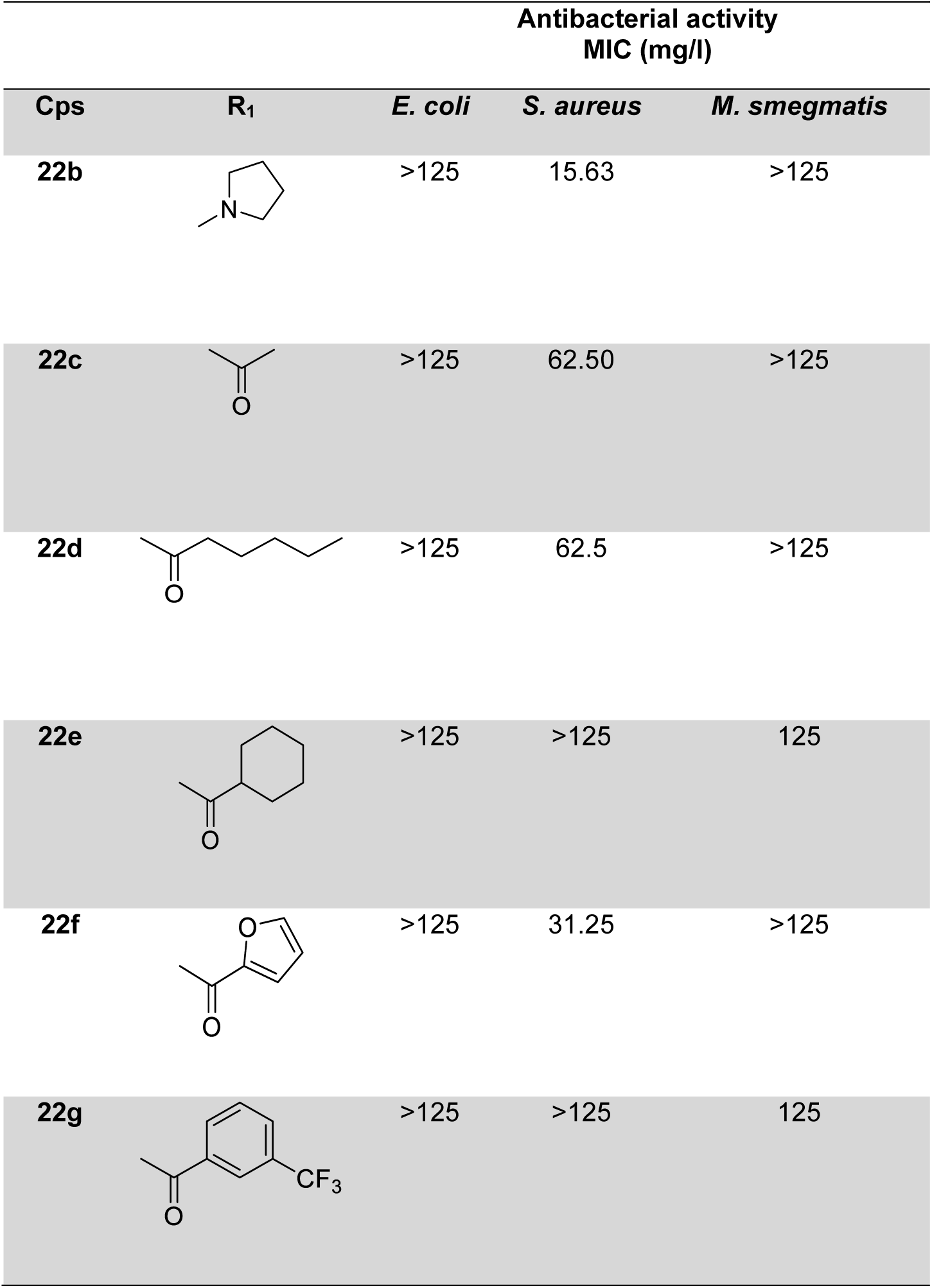
Antibacterial activities of the C-6 derivatives. (R_1_: residue on C-6 position of quinoxaline core)

**Figure 4.**
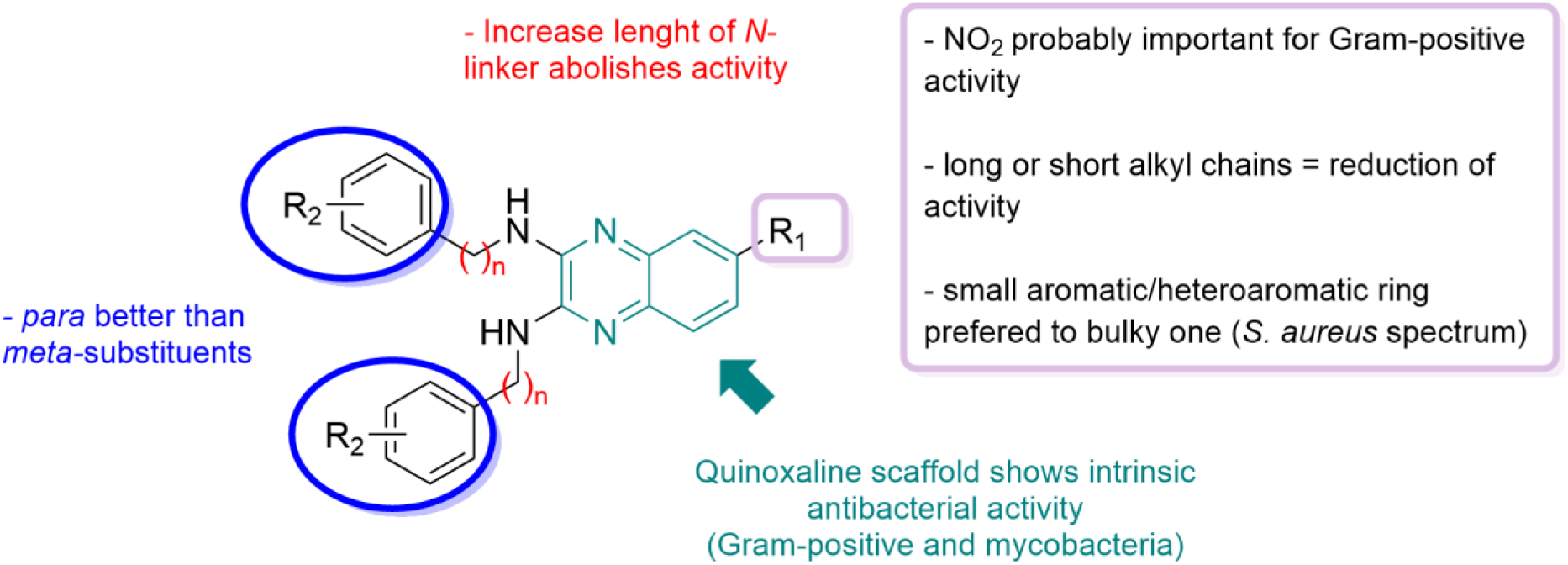
Summary of the SAR studies performed on different regions of the synthesised derivatives. A total of 15 similarly structured compounds were analysed to generate this map. All the biological results regarding their antibacterial activity were included in **Tables 1-2**.

### Discussion around Phase 2 and 4 of this study

A sub-selection of these quinoxaline derivatives (namely compound **22f**, **25**, **31**, **32**, **35** and 2,3-dichloro-6-nitroquinoxaline/**2Cl-Q**) was selected for additional testing (Phase 2) against 19 clinically relevant strains as they showed initial activity against MRSA (**Figure 1**).

Here, we confirmed the importance of the *N*-aromatic ring’s functionalisation for the antibacterial activity against *S. aureus* due to negligible activity of the central scaffold (**2Cl-Q**) and of the C6 derivative (**22f**). Compounds **25**, **31**, **32** and **35** retained activities against the *S. aureus* clinical isolates (confirming Phase 1’s results) and were even more active against the documented drug-resistant strains (**Table S2B**). This was particularly noticeable for compounds **25** and **31**.

We also obtained data about the antibacterial activity of the quinoxaline derivatives against *E. faecalis*. Similarly to *S. aureus*, compounds **25**, **31**, **32** and **35** were very active against both clinical and drug resistant isolates including the most important *vanA* and *vanB* genotypes (**Table 3**). More specifically, *S. aureus* and *E. faecalis* expressing clinical resistance to systemic antibiotics (flucloxacillin, vancomycin, tetracycline and erythromycin/clindamycin) were both found to be susceptible to compounds **25** and **31** at low concentrations (0.06-0.25 mg/l - **Table 3**).

We extended our testing against a larger spectrum of Gram-positive bacteria (**Figure 2**) including *S. aureus*, *E. faecalis* and *E. faecium* to ensure our selected quinoxaline derivatives (namely compounds **25** and **31**) retained their activity against a broad range of clinical isolates including some having specific and documented antibiotic resistance mechanisms (**Table S3**). *E. faecium* was of particular interest due to its broader resistance and higher virulence compared to *E. faecalis* [30].

Encouragingly, compounds **25** and **31** maintained potent antibacterial activity against a wide panel of MRSA strains (**Table S3**), including clinically important pathogens with reduced clindamycin susceptibility (via the D-zone test [36]) and containing Panton-Valentine Leukocidin (PVL) genes (*pvl* genes detected by PCR [37]). They demonstrated better antibacterial profiles than vancomycin and teicoplanin (first line treatment) and linezolid (second line treatment) with MICs ranging from 0.125 to 0.500 mg/l and comparable activity to the cyclic lipopeptide antibiotic daptomycin (**Figure 2**). In terms of *E. faecalis* and *E. faecium*, compounds **25** and **31** showed equal to or better activity with lower concentration than most of the antibiotics tested except for teicoplanin and daptomycin against *E. faecalis* 23948 and vancomycin against *E. faecium* 26575. These results are particularly promising when considering the impact that both *Enterococci* have on urinary tract infections and hospital length of stay (i.e., *E. faecium*) [38] as well as sepsis, endocarditis and meningitis in immunocompromised patients (i.e., *E. faecalis*) [39, 40].

One of the main challenges in developing new antimicrobials relates to how long the targeted bacterial species would need to develop resistance [1, 2]. Therefore, we decided to assess how pre-exposure to our quinoxaline derivatives (namely compounds **25** and **31**) affects bacterial susceptibility. Pre-exposure of all tested bacterial strains to sub-inhibitory doses of compounds **25** and **31** led to increases in MIC levels (**Table 4**). The increases were more profound for pre-exposure to compound **31**. Yet pre-exposure to compounds **25** and **31** did not alter the clinical susceptibility profile of commonly used antibiotics, with the expectation of trimethoprim/sulfamethoxazole and vancomycin in one of the two *E. faecalis* strain tested (*E. faecalis* NCTC 12201 - **Table 5**).

The pre-treatment of vancomycin resistant *E. faecalis* (NCTC 12201) with compound **31** (but not compound **25**) made this strain sensitive to vancomycin. At this stage we can only speculate about the potential mechanism of resistance involved. Efflux, which is a common and efficient mechanism leading to multidrug resistance [41], is likely to be involved here (no change in tetracycline or quinolones (levofloxacin and ciprofloxacin) MIC – **Table 5**). Compound **31** might inhibit efflux pumps critical for expelling a wide range of structurally diverse compounds [42] in vancomycin resistant *E. faecalis* (NCTC 12201). A recent study also demonstrated that structurally-similar quinoxaline-containing compounds inhibited efflux pump activity and restored drug susceptibility in drug-resistant non-tuberculous mycobacteria [43].

In contrast, pre-treatment of *E. faecalis* (NCTC 12201) with compound **31** induced a sensitive to resistant transition to trimethoprim/sulfamethoxazole (**Table 5**). Once again, the mechanism behind this transition is currently unknown. Despite this, our lead compounds **25** and **31** were quite active against multidrug resistant pathogens (**Figure 1** and **Table 3**).

While MIC values represent a measure of antibacterial susceptibility (**Tables 3** and **Table 4**), they do not reveal whether the antibiotic (or test compound) is bacteriostatic or bactericidal [44]. Recent studies support the combined use of MIC and MBC to provide a more detailed understanding of the bacteria susceptibility to compounds [45] and correlate *in vitro* data with possible outcomes of *in vivo* treatments [46]. In our investigations, the MBC of compounds **25** and **32** (**Table 6**) were higher than their MIC values (**Table 3** and **Table 4**) confirming the general trend of MBC being higher or equal to MIC [47]. Moreover, we can conclude that compound **25** is bactericidal since the MBC is no more than four times the MIC value [48, 49].

In addition to quantifying compound **25** and **31**’s MICs and MBCs on *S. aureus* and *E. faecalis*, we also measured these quinoxaline-containing compounds’ ability to affect biofilms. Biofilms have been recognised as a potential source of recurring infection and high levels of antibiotic tolerance are prevalent in bacterial biofilms [50]. Equally, the formation of biofilms on implant surfaces is a major cause of implant-associated infection difficult to treat [51]. Biofilm-producing bacteria show a different behaviour when compared to planktonic (free-floating) bacteria that are typically used in the testing of traditional antibiotics; this behaviour often limits compound penetration through biofilms [52]. Reassuringly, both compounds **25** and **31** retained their antibacterial against pre-formed *S. aureus* and *E. faecalis* biofilms and, more importantly, they performed better than all other antibiotics tested (except for rifampicin for *S. aureus* biofilm – **Table 7**). This result is particularly promising for the use of these compounds in antibiofilm products like implants and wound dressings, although further testing would be required against biofilms *in vivo*, which are notably more resistant to antibiotics than *in vitro* ones [51].

## Conclusion

This study identified compounds with promising antibacterial activities, especially against a collection of clinical isolates of Gram-positive bacteria including *S. aureus*, *E. faecalis* and *faecium*. The identification of quinoxaline derivatives (particularly compounds **25** and **31**) with comparable (or better) activity to currently used antibiotics (vancomycin, teicoplanin, linezolid and daptomycin) expands the portfolio of chemical matter to be further developed for clinically relevant pathogens [2].

Furthermore, we also showed that compounds **25** and **31** had efficacy against bacterial biofilms, which is an important property rarely tested at this stage of investigation [51].

Further medicinal chemistry optimisations will allow a more detailed examination of the SAR around these compounds leading to the identification of even more potent antibacterial drug candidates with biocidal and biofilm-inhibiting/eradicating characteristics. At present, due to the poor aqueous solubility of these two compounds, they are most likely to be used in topical formulations for treatment of uncomplicated skin and soft-tissue infections or in the coating of implants or dressings.

## Methods

### Workflow of antibacterial investigations described in this study

The present study was performed in four phases. In Phase 1, a library of 15 compounds (13 synthesised compounds and the 2 initial building blocks) was initially tested against nine bacterial strains (see full list in **Table S1**). In Phase 2, only the six most promising compounds derived from Phase 1 were investigated against 18 bacterial species originating from clinical settings. In Phase 3, the antibacterial activities of the two most active compounds (derived from Phase 2) were next explored against a further 41 clinical strains. In Phase 4, the effects of pre-exposure to sub-minimum inhibitory concentrations (MIC) were evaluated of the two most active compounds (derived from Phase 2).

The organisms used in this study (full list included in **Table S1**) were obtained from hospital Laboratories (e.g., SACU_Bead_No 23306, available from the Specialist Antibacterial Chemotherapy Unit - Public Health Wales, Cardiff), the American Type Culture Collection (e.g., ATCC 700699, 12301 Parklawn Drive, Rockville, MD 20852, USA) and the National Collection of Type Cultures (e.g., NCTC 12201, Central Public Health Laboratory, 61 Colindale Avenue, London NW9 5HT).

### Phase 1: Compound preparation

A library of 15 compounds (13 synthesised compounds and the 2 initial building blocks, described in Padalino *et al*. [11]) was prepared in 100% methanol at 2.5 mg/ml final concentration. A full list of compounds can be found in **Table S1**.

### Phase 1: bacterial growth conditions

All procedures were carried out in a biosafety level (BSL) 2 cabinet. A fresh subculture of each of the nine bacterial species (full list in **Table S1**) was prepared by streaking onto a fresh agar plate and incubating at 37°C for 24 h. The agar plates were prepared with high salt Luria-Bertani medium (HSLB, 4 g), agar (2 g) and water (200 ml) for all strains except *Mycobacterium smegmatis* mc^2^15, which required supplementation of the solid growth medium with 0.2% v/v glycerol and 0.05% v/v Tween-80. Bacteria were stored on agar plates at 4°C until needed and replaced weekly.

Prior to use, a single colony of each bacterium was removed from the agar plates using a sterile loop and inoculated in fresh growth medium. Luria-Bertani (LB) medium was used for all strains except *M. smegmatis* mc^2^15, which required supplementation with 0.2% (v/v) glycerol and 0.05% (v/v) Tween-80. Cultures were incubated for 24 h (or 48 h for *M. smegmatis*) at 37 °C with aeration at 200 rpm until they reached an OD_600_ between 0.8 and 1.0 (assessed using spectrophotometer BioTek Synergy 4). In the case of low OD_600_, the cultures were left to incubate further; if the OD_600_ was higher than 1.0, then a dilution was performed. Once optimal OD_600_ were reached, each bacterial culture was diluted with LB medium to approximately 1.0 × 10^5^ CFU/ml.

### Phase 1: Determination of *in vitro* antibacterial activity against bacteria isolates

The MIC was determined using the broth microdilution method in a 96-well plate containing fresh LB medium except for *M. smegmatis*, which was supplemented with 0.05% Tween 80 and 0.2% glycerol [19–21]. Full list of bacteria isolates included in **Table S1**.

A primary screen was carried out at both 125.0 mg/l and 62.5 mg/l to keep the MeOH content below 10% v/v. A secondary, dose response titration (125.00, 62.50, 31.25, 15.63, 7.81, 3.91 mg/l or even lower concentrations when appropriate) was performed only for compounds able to inhibit the visible growth of bacteria in the primary screen. In each assay, all compounds were tested in triplicate against the nine bacteria strains; both primary and dose response assays were performed twice.

The OD_600_ was measured at the beginning (initial reading) and at the end (after incubation at 37°C for 24 h or 72 h for *M. smegmatis* – final readings) of dose response titration. Those readings were compared to calculate the MIC (as mg/l), defined as the lowest concentration of compound that inhibits 90% of the growth of the organism studied.

### Phase 2 and 3: determination of *in vitro* antibacterial activity against clinically relevant bacteria strains

A sub-selection of compounds (**2Cl-Q**, **22f**, **25**, **31**, **32** and **35**) was selected for Phase 2 of the study (**Table S2**). Here, the broth microdilution assay was performed in Mueller-Hinton broth (MHB) according to the ISO-20766 international standard [22] and clinical significance of MIC was interpreted using the current EUCAST breakpoints (https://www.eucast.org/clinical_breakpoints/).

Each compound was prepared in DMSO (instead of MeOH used in Phase 1) and then diluted in water to create stock solutions at lower concentrations (0.008mg/l to 128mg/l). For some bacteria (*S. pneumoniae*, *H. influenzae* and *Neisseria* species), the MHB was supplemented with 5% lysed horse blood and nicotinamide adenine dinucleotide (β-NAD).

The MIC values (expressed in mg/l) were determined as the lowest concentration that, under defined *in vitro* conditions (incubation at 34 °C to 37°C), prevented visible growth of bacteria within a defined period of time (for 18-24 h).

During Phase 3, four antibiotics (vancomycin, teicoplanin, linezolid and daptomycin) with known activity against Gram-positive agents were used for activity comparison (**Table S3A**). The MIC values (expressed as mg/l) of each known antibiotic were interpreted using the current EUCAST breakpoints [22] (https://www.eucast.org/clinical_breakpoints/ - summarised in **Table S3B**).

### Phase 4

In this final stage of the study, only the two most active compounds (**25** and **31**) were further investigated to determine potential bacterial emerging resistance following exposure, their minimum biocidal concentrations (MBCs) and their minimum biofilm eradication concentrations (MBECs).

### Phase 4: Determination of MIC following pre-exposure to quinoxaline analogues 25 and 31

Fresh stocks of compounds **25** and **31** were resuspended in 1 ml DMSO and further diluted to 512 mg/l in deionised water (final DMSO concentration below 1% v/v). *S. aureus* ATCC 29213, *E. faecalis* ATCC 29212 (vancomycin sensitive) and *E. faecalis* NCTC 12201 (vancomycin resistant) (see details in **Table S1**) were stored in cryopreservation beads (Fisher Scientific, Loughborough, UK) at −80°C and sub-cultured onto tryptone soya agar (TSA) for a maximum of two subcultures before use.

To initiate liquid cultures of each strain for repeat MIC determinations, MHB supplemented with cations to a final concentration of 20 mg/l CaCl_2_ and 10 mg/l MgCl_2_ was inoculated with 2-3 bacterial colonies and incubated at 37°C for 16-24 h. Suspensions were centrifuged at 3,000 x g for 20 min to pellet the bacteria; the pellets were resuspended in fresh cation-adjusted MHB to reach a cell density between 1.5 – 5 ×10^6^ CFU - Colony Forming Units/ml.

Following the ISO-20766 international standard [22], assays were initiated in 96-well microtiter plates where descending 2-fold dilutions of compounds (quinoxaline analogues or antibiotic controls) were included in a final volume of 50 μl; negative control wells remained compound free but contained 50 μl of MHB instead. Aliquots of adjusted bacterial inoculum (50 μl) were added to each well except for the negative control wells where 50 μl of MHB was added instead, giving a final cell concentration of approximately 5 × 10^5^ CFU/ml. The plates were incubated at 37°C for 16-24 h. The lowest concentration of compound that inhibited cell growth was determined by visual inspection and recorded as the MIC (mg/l). The experiments were conducted in triplicate and the most frequently occurring MIC was recorded.

To determine MIC following compound pre-exposure, bacterial cultures were exposed to compounds **25** and **31** (at half of the MIC concentration, determined above) for a period of 16-24 h at 37°C in total volumes of 10 ml in 50 ml Falcon™ centrifuge tubes (Fisher Scientific, UK), using the cell densities and culture conditions described in the MIC protocol. Bacterial cultures were incubated with agitation at 200 rpm. Following incubation, the cultures were centrifuged at 3,000 x g and the pellets were resuspended in neutraliser solution (Lecithin 10 g/L, Tween80 30 g/L, Sodium Thiosulphate 20 g/L, L-Histidine 1 g/L, Saponin 30 g/L, Sodium Dodecyl Sulphate 5 g/L in deionised water) and vortexed. The suspensions were re-centrifuged, the cell pellets resuspended in tryptone sodium chloride (TSC) and the cell densities adjusted to 1.5 - 5 × 10^6^ CFU/ml.

MIC determination of compounds **25** and **31** were subsequently determined as described above using the cultures that had been pre-exposed to sub-MIC levels of the test compounds.

In addition, a selection of known antibiotics (ampicillin, imipenem, vancomycin, levofloxacin, ciprofloxacin, trimethoprim/sulfamethoxazole, cefotixin, gentamicin, erythromycin, tetracycline, rifampicin and benzylpenicillin) was diluted and prepared as described in BS EN ISO20776-1:2020 [26]. Antibiotic loaded 6 mm paper disks (Oxoid, Bassingstoke, UK) were applied to the surfaces of the agar plates seeded with each of the three bacteria strains (stated above) before (control) and after pre-exposure to test compounds **25** and **31**. Plates were incubated for 18 ± 2h at 37°C. Zones of inhibition were measured using a calliper and recorded. The antibiotic susceptibility profiles of the three bacteria strains before and after pre-exposure to the quinoxaline derivatives to the known antibiotics selection were compared to the current EUCAST breakpoints (https://www.eucast.org/clinical_breakpoints/) [27].

### Phase 4: Determination of minimum biocidal concentration (MBC)

Following MIC determination (described in the section above), the entire contents of the wells corresponding to the MIC level and all higher concentrations for which no visible growth was observed, were removed by pipette and plated onto TSA plates. The plates were incubated at 37°C for 16-24 h. The MBC was recorded as the lowest concentration of test compounds for which there was no colonies.

### Phase 4: Determination of minimum biofilm eradication concentration (MBEC)

The standard ASTM E2799 [28] assay was used to determine the MBEC of compounds **25** and **31** as well as several antibiotics (vancomycin, rifampicin, linezolid, teicoplanin and sparfloxacin). Briefly, bacterial cultures (1.5 × 10^5^ CFU/ml) were added to 96-well Calgary biofilm devices (Innovotech, Canada); plastic lids containing 96 pegs were subsequently added to each 96-well base. The entire Calgary device was incubated at 37°C for 16-24 h in an orbital shaker to allow the initial biofilm establishment on a surface. The biofilm-containing pegs were rinsed in TSC to remove planktonic cells and transferred to a challenge plate containing test compounds **25** and **31** as well as five selected antibiotics serially diluted across the plate. Following another 24 h incubation, the pegs were placed in neutralising broth (Lecithin 10 g/L, Tween-80 30 g/L, Sodium Thiosulphate 20 g/L, L-Histidine 1 g/L, Saponin 30 g/L, Sodium Dodecyl Sulphate 5 g/L in deionised water) for 10 min and then transferred to a recovery plate containing 100 μl of sterile tryptone soya broth (TSB). An effectiveness test of the neutralising broth was performed in accordance with ASTM E2799 to validate use of the neutralising broth. The recovery plates were placed in a sonicating water bath for 30 ± 5 mins to disaggregate the biofilms. The lids containing the pegs were discarded and replaced with standard lids. The plates were incubated for 37°C for 16-24 h to allow biofilm growth and the MBECs were determined qualitatively by recording the lowest concentration of antibiotic/compound that prevented cell growth (i.e. absence of turbidity).

### Evaluation of cell morphology following exposure to test compounds

Bacterial cultures (*S. aureus* ATCC 29213 and *E. faecalis* NCTC 12201) were grown for 16-24 h at 37°C in cation adjusted MHB and resuspended in TSC to approximately 1 × 10^9^ CFU/ml.

A suspension of bacteria (final concentration 5 × 10^5^ CFU/ml) and each compound (**25** and **31**) was prepared at a concentration being double the MBC of each test compound.

Untreated control cultures were initiated using TSC only. Following incubation at 37°C for 16-24 h, the suspensions were centrifuged at 3,000 x *g* and bacteria pellets fixed by incubation with 2.5 % glutaraldehyde for 2 h at 21°C. Following another centrifugation step, the cell pellets were washed in an ascending series of ethanol concentrations (from 10 and 100 %), with 5 min incubation at each step and centrifugation between each wash. The entire 2 ml volume of the final suspension in 100 % ethanol was filtered through a 0.2 μm polycarbonate membrane (Whatman™, Cytiva, UK) using a manifold system. The filtered membranes were transferred to petri dishes and placed in a bell-jar overnight to remove residual moisture.

Membranes were subsequently fixed to aluminium stubs using carbon adhesive tabs (Fisher Scientific, Loughborough, UK) and sputter-coated with 20 nm gold/palladium (Au/Pd). Scanning electron microscopy (SEM) images were acquired using a beam energy of 5kV and an in-lens detector on a Sigma HD field gun Scanning Electron Microscope (Carl Zeiss Ltd, UK). Three representative fields of view were captured for each treatment at magnifications between 10,000 and 50,000.

## Acknowledgment

The authors thank Dr Sumana Bhowmick for the initial training on *in vitro* antibacterial screening at the beginning of Phase 1 of the study as well as Dr Mandy Wootton and Ms Jennifer Richards from the Specialist Antibacterial Chemotherapy Unit (Public Health Wales, Cardiff) for having conducted the antibacterial screens during Phase 2 and 3 of this study. This work was supported by Aberystwyth University (Technology Transfer Grant Development Award) and the Life Sciences Wales Research Network (a Welsh Government Ser Cymru initiative).

## Notes

### Competing Interest Statement

The authors have declared no competing interest.

